# Compensatory evolution facilitates loss of *prfB* autoregulation in *Pseudomonas fluorescens* SBW25

**DOI:** 10.64898/2026.01.08.698412

**Authors:** Sungbin Lim, Frederic Bertels, Javier Lopez-Garrido, Jenna Gallie

## Abstract

Understanding why some traits are maintained whereas others are repeatedly lost is a central question in evolutionary biology. Here, we address this problem through an analysis of the evolutionary dynamics of autoregulation of the *prfB* gene, which encodes peptide-chain release factor 2 (RF2), a key factor in bacterial translation termination. RF2 recognizes UGA and UAA stop codons and catalyzes the release of the completed polypeptide. In many species, *prfB* contains an internal UGA stop codon, triggering premature translation termination by RF2 itself. Complete RF2 translation depends on a +1 programmed ribosomal frameshifting (PRF) event on the internal stop codon, which occurs more frequently when RF2 levels are low, resulting in autoregulation of *prfB* expression. While widespread, this autoregulatory mechanism has been lost in multiple bacterial lineages. We combined phylogenetics, experimental evolution and molecular genetics to investigate the evolutionary forces behind this loss. We found no significant correlation between PRF loss and stop codon usage using phylogenetically informed analyses, and PRF disruption in *Pseudomonas fluorescens* SBW25 had no detectable fitness effect. However, engineered mutations that reduced frameshifting caused fitness defects, which were compensated by two classes of mutation: (i) mutations that impair specific ribosomal proteins, and (ii) single-nucleotide deletions in *prfB* that adjust the reading frame and bypass the internal stop codon. These results suggest that compensatory mutations facilitate the loss of *prfB* autoregulation under RF2-limiting conditions. We discuss three potential scenarios that could account for this process.

## INTRODUCTION

Translation fidelity is essential for cellular homeostasis, and a critical step is the accurate recognition of stop codons during termination. This process relies on ribosome release factors (RFs), which bind to stop codons and catalyze the hydrolysis of the peptide bond between the nascent polypeptide and the tRNA in the peptidyl (P) site, thereby releasing the complete protein and promoting ribosome disassembly. While a single RF recognizes all three stop codons in eukaryotic cells, bacteria possess two RFs: RF1, which recognizes UAG and UAA, and RF2, which recognizes UGA and UAA.

Although precise recognition of stop codons by RFs is critical for translation fidelity, not all stop codons are equally efficient (Adamski, Donly and Tate 1993; Poole, Brown and Tate 1995; Pavlov *et al*. 1998; Higashi *et al*. 2006; Betney *et al*. 2010; Wei and Xia 2017; Zhang *et al*. 2020; Biziaev *et al*. 2022; Riegger and Caliskan 2022; Loughran *et al*. 2023). Some are prone to be bypassed at high rates due to ribosomal frameshifting (Brown *et al*. 1993, 1994), and in certain cases these inefficient stop codons have been co-opted for gene regulation or the production of alternative protein isoforms (Wong *et al*. 2008; Antonov *et al*. 2013; Caliskan *et al*. 2017; Fan *et al*. 2017). A well-documented example is the autoregulation of *prfB*, the gene encoding RF2, which contains a highly inefficient UGA stop codon within its reading frame (Craigen *et al*. 1985; Craigen and Caskey 1986; Weiss *et al*. 1988) (Figure 1A). Recognition of the internal stop by RF2 results in translation termination after only 24 residues in *E. coli*, yielding an unstable peptide that is rapidly degraded (Williams *et al*. 1989). Alternatively, the internal stop can be bypassed by a +1 programmed ribosomal frameshift (PRF), allowing the synthesis of full-length RF2 (Donly *et al*. 1990; Mikuni, Kawakami and Nakamura 1991). The balance between termination and frameshifting is affected by RF2 levels: low levels increase frameshifting frequency and the production of full-length RF2, whereas high levels promote termination, creating a negative feedback loop (Figure 1A).

**Figure 1.**
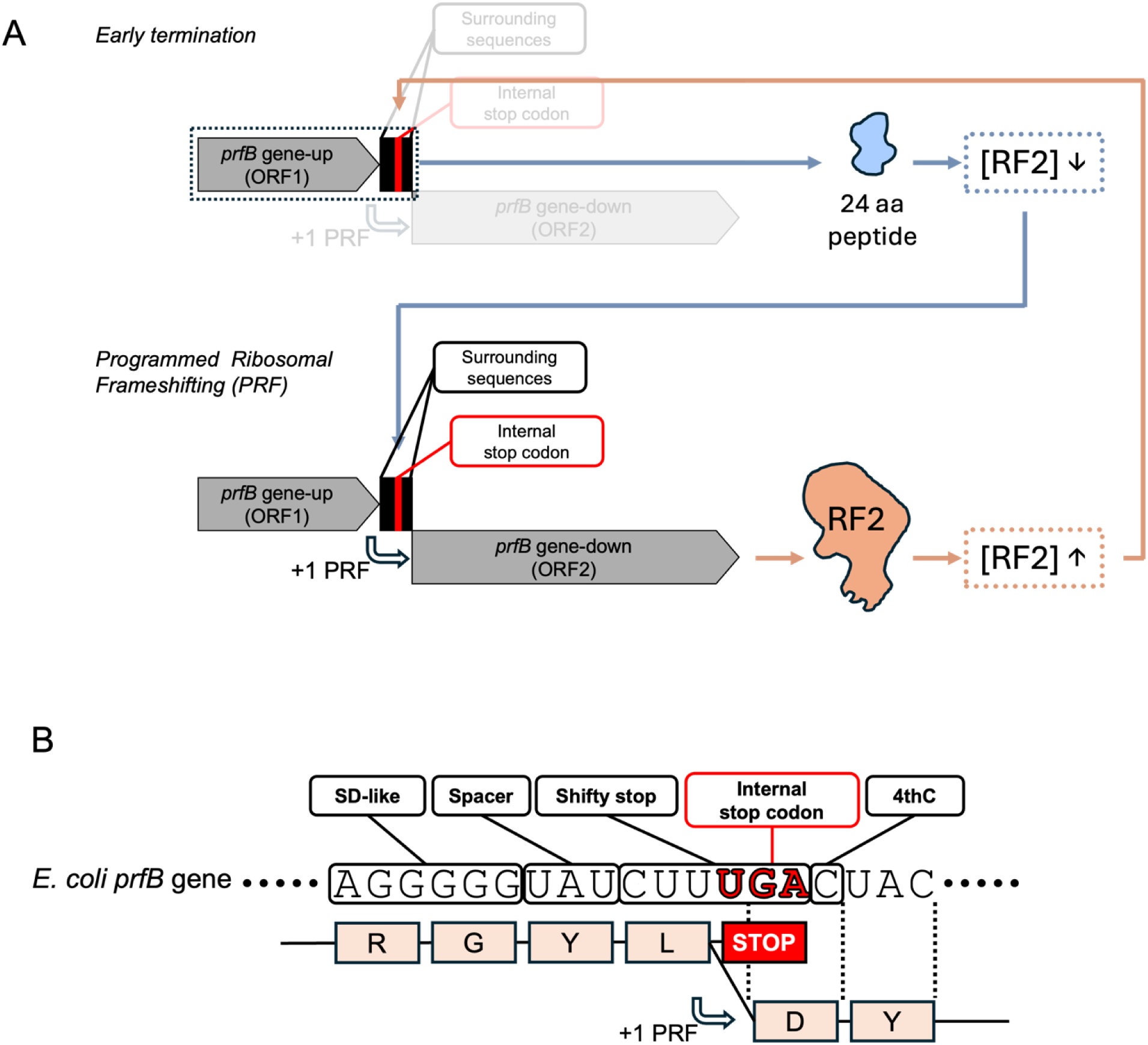
Programmed Ribosomal Frameshifting (PRF) in *prfB*. (**A**) Diagram of *prfB* autoregulation. The *prfB* locus is depicted as two gene arrows—*prfB* gene-up (ORF1) and *prfB* gene-down (ORF2)—separated by an internal UGA stop codon (red box). Black boxes flanking the UGA indicate sequences required for frameshifting (see panel B for details). Synthesis of full-length RF2 requires a +1 programmed ribosomal frameshift (+1 PRF) that bypasses termination at the internal UGA. When RF2 concentration is high (upper panel, early termination), the internal UGA is efficiently recognized, causing premature termination and production of an unstable 24-aa peptide (blue). When RF2 concentration is low (lower panel, PRF), the +1 frameshift occurs at high frequency, bypassing the internal stop and yielding full-length RF2 (orange), thereby increasing RF2 levels. This autoregulatory loop maintains RF2 homeostasis. The orange line indicates the functional path in which increased frameshifting raises RF2 concentration; the blue line shows the opposite scenario. (**B**) Diagram showing functionally important elements in the PRF site of *E. coli prfB* gene. The UGA stop codon is in red. Additional elements that promote frameshifting are indicated: a Shine–Dalgarno (SD)-like sequence (AGGGGG), a UAU spacer codon, the shifty-stop sequence (CUU UGA) and the C at the nucleotide immediately following the UGA stop codon (4thC). The amino acids produced by translation before and after the +1 frameshift (+1 PRF) are indicated below the RNA sequence. See main text for details.

The mechanics of the PRF event have been best characterized in *E. coli*. The PRF site spans 16 nucleotides and includes several elements that influence frameshifting efficiency (Figure 1B) (throughout the manuscript, nucleotide annotations at PRF site are given in RNA notation, using U instead of T). The most critical component is the “shifty stop” sequence (CUU UGA), which favors +1 frameshifting via wobble base pairing between tRNA-Leu(GAG) and the mRNA (Weiss *et al*. 1987, 1988; Curran and Yarus 1988). Briefly, the CUU codon is decoded by wobble tRNA-Leu(GAG); if the anticodon shifts +1, it can still pair with the new codon UUU. This shift is enhanced by a nearby Shine-Dalgarno (SD)-like sequence (AGG GGG), which transiently stalls the ribosome when the CUU codon is in the P-site, thereby facilitating frameshifting (Weiss *et al*. 1987, 1988). The spacer codon immediately preceding the "shifty stop" sequence, typically UAU or UCU (Baranov, Gesteland and Atkins 2002a), also promotes frameshifting by favoring rapid tRNA ejection (Baranov, Gesteland and Atkins 2004; Devaraj *et al*. 2009). tRNAs with stronger base pairings slow ejection, and reduce frameshifting (Márquez *et al*. 2004; Sanders and Curran 2007). Finally, the nucleotide immediately after the stop codon—hereafter referred to as the 4^th^ position—influences termination efficiency, with cytosine particularly enhancing frameshifting (Poole, Brown and Tate 1995). Together, these features create a tuneable regulatory site in which translation termination competes with +1 frameshifting to control RF2 synthesis.

Nearly all naturally occurring bacteria encode RF2, and at least two thirds have an internal stop codon in *prfB* (Persson and Atkins 1998; Baranov, Gesteland and Atkins 2002a; Bekaert, Atkins and Baranov 2006; Naeem *et al*. 2023). The PRF-associated elements tend to be highly conserved across species (Baranov, Gesteland and Atkins 2002a, 2002b; Prince, Lin and Feaga 2025), and rare variability in this region has been correlated with changes in the frameshifting rates (Craigen and Caskey 1986; Weiss *et al*. 1988; Naeem *et al*. 2023). Despite this widespread conservation, the physiological relevance of *prfB* autoregulation remains unclear. Autoregulation is often proposed to prevent RF2 overproduction (Betney *et al*. 2010; de Silva *et al*. 2010), which may help minimize mis-termination at sense codons similar to UGA (Freistroffer *et al*. 2000; Prince, Lin and Feaga 2025), such as the tryptophan codon UGG (Abdalaal *et al*. 2020), or to regulate the frequency of ribosome drop-off during entry to stationary phase (Betney *et al*. 2010). However, studies in different species have found that deletion of the internal stop has subtle or negligible deleterious effects under laboratory conditions (Johnson *et al*. 2011; Abdalaal *et al*. 2020; McNutt *et al*. 2021; Naeem *et al*. 2023), and only *prfB* overexpression from inducible promoters have been associated with growth defects (Jørgensen *et al*. 1993; Rengby and Arnér 2007; Lalanne, Parker and Li 2021; Prince, Lin and Feaga 2025). In addition, published surveys indicate that autoregulation is completely absent from as many as one third of sequenced genomes (Persson and Atkins 1998; Baranov, Gesteland and Atkins 2002a; Prince, Lin and Feaga 2025), suggesting that the mechanism can be lost without fatal fitness consequences in nature.

Here, we have explored the evolutionary forces behind *prfB* autoregulation loss using *P. fluorescens* SBW25 as a model system. As in other species, elimination of *prfB* autoregulation had no detectable fitness effect under standard laboratory conditions. However, mutations in the PRF site that reduced frameshifting efficiency led to severe fitness defects. Through experimental evolution, we identified two compensatory strategies that alleviate these defects: knockdown of ribosome-associated proteins (RsmA, RsmH and RplI) that globally reduce decoding fidelity, and single-nucleotide deletions upstream of the internal stop codon that abolish autoregulation. We propose that the evolutionary dynamics of the internal stop codon hinge on its extreme tunability. Mutations in the PRF site tend to be deleterious, placing organisms on a sharp fitness peak where most mutations are harmful. In this view, loss of the internal stop codon may represent an evolutionary escape from this constraint.

## RESULTS

### Characterization of *prfB* autoregulation in *P. fluorescens*

The *prfB* gene of *P. fluorescens* carries an internal stop after 24 codons, flanked by canonical PRF site elements: a SD-like sequence (CGG GGG), separated by a UAU spacer codon from a shifty CUU UGA stop, and a cytosine at the 4th position, immediately downstream of the stop codon (Figure 2A). These elements likely determine frameshifting rate on PRF, through the mechanism illustrated in Figure 1A.

**Figure 2.**
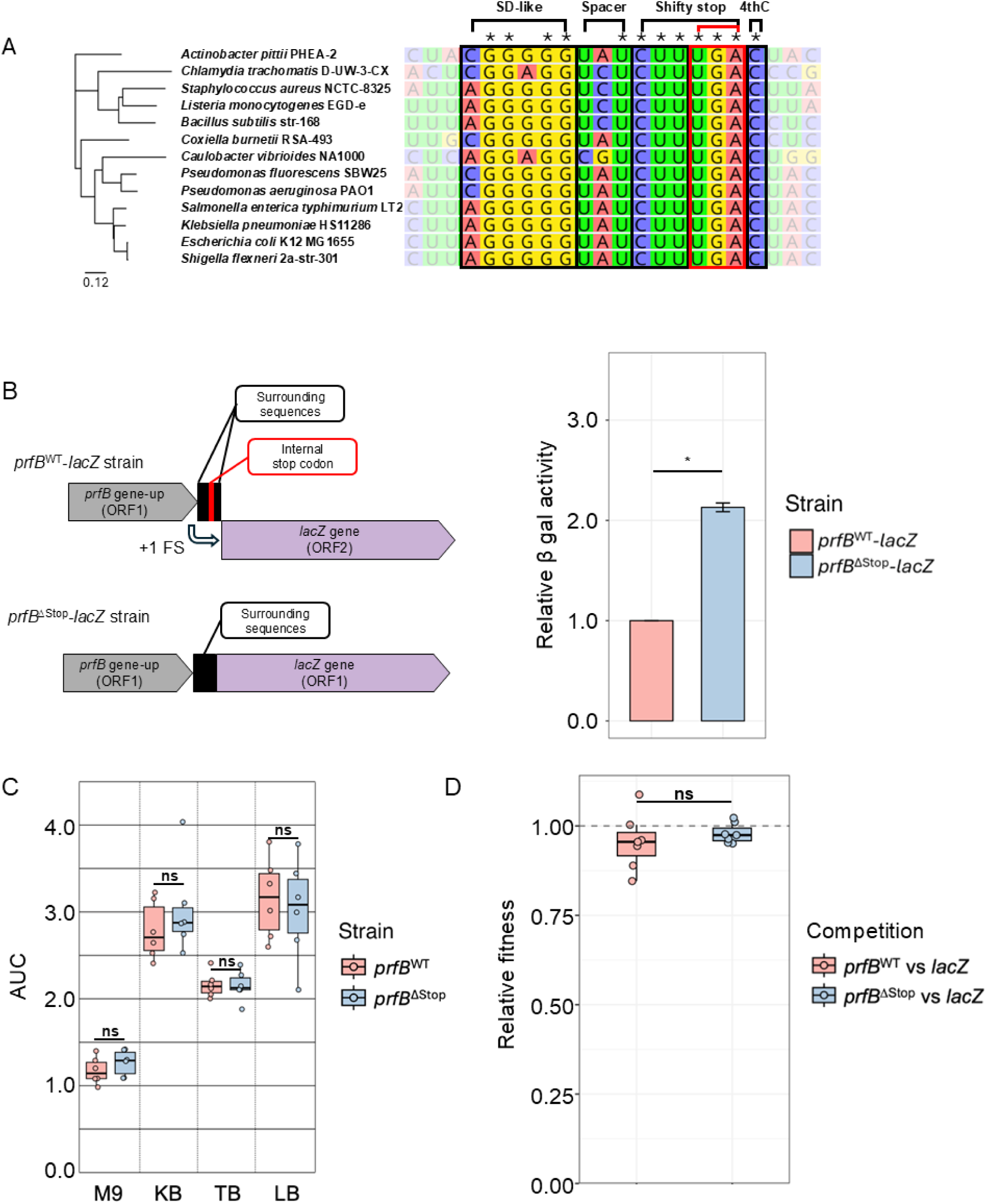
Characterization of PRF in *P. fluorescens prfB.* (**A**) Nucleotide alignment of PRF sites from 13 bacterial species. Asterisks mark positions conserved across all sequences. The species tree (left) was inferred from 16S rRNA; the scale bar indicates branch length. Boxes highlight PRF-site features: black boxes denote flanking elements required for frameshifting (SD-like sequence, spacer codon, 4thC and shifty-stop sequence) and red boxes denote UGA stop codon. (**B**) β-galactosidase activity of *prfB*^WT^-*lacZ* and *prfB*^ΔStop^-*lacZ* fusions. Left, schematics of the fusions: the *prfB* upstream fragment extending to the PRF site was placed upstream of *lacZ*, either retaining (WT) or removing (ΔStop) the first nucleotide of the internal UGA stop codon. Right, β-galactosidase activity was expressed relative to the wild type (*prfB*^WT^-*lacZ*). Data represent the average and standard error of five independent experiments. (**C**) Area under the growth curve (AUC) for *P. fluorescens* strains carrying the wild-type *prfB* allele (*prfB*^WT^, orange) or a mutant lacking the internal stop codon (*prfB*^ΔStop^, blue). Cultures were grown in M9 minimal medium with 0.4% (w/v) Glucose (M9), King’s Broth (KB), Terrific Broth (TB), or Lysogeny Broth (LB). The value of each condition was represented as box plots. A horizontal line in the middle of the box indicates the median if the number of replicates is odd. If the number of replicates is even, the central two values are averaged and plotted as a horizontal line. The box covers the interquartile region (IQR) of each dataset, while the whiskers reach to the extreme individual value within 1.5 times the IQR from the median value. Each biological replicate (six in this case) is shown as an individual circle. This formatting on the box plot stays consistent across other box plots in the manuscript, while the number of replicates varies on each figure. (**D**) Competition assays of *P. fluorescens* strains carrying the wild-type *prfB* allele (*prfB*^WT^, orange) or the *prfB*^ΔStop^ allele (blue) against an isogenic reference strain bearing wild-type *prfB* and constitutive *lacZ* (labelled as *lacZ*) for colony discrimination on plates with X-gal. Assays were performed in King’s Broth (KB). A relative fitness of 1 is indicated by the dashed line. Individual data points are shown as circles (n = 7 per group). ‘*’ p ≤ 0.05; ‘ ns’ p > 0.05.

To experimentally test the functionality of *P. fluorescens* PRF, we deleted the first nucleotide of the internal UGA stop codon (ΔStop), restoring the reading frame and bypassing the need for frameshifting to produce RF2. We quantified the impact of the deletion in RF2 production using two approaches. First, we generated translational *prfB-lacZ* fusions, in which *lacZ* was placed immediately after the 4^th^ position, following the wild type stop codon (*prfB*^WT^-*lacZ*), or the ΔStop allele (*prfB*^ΔStop^-*lacZ*). Both constructions were constitutively expressed and inserted at a neutral ectopic site in the *P. fluorescens* chromosome, between loci *pflu1179* and *pflu1180* (Zhang and Rainey 2007). β-galactosidase activity measurements indicated that the lack of stop codon increased expression by more than twofold (Figure 2B). Second, we raised an anti-RF2 antibody and directly measured the levels of RF2 in the wild type and in the *prfB*^ΔStop^ strain (Supplementary Figure S1). The deletion of the internal stop codon resulted in a clear increase in RF2 levels, demonstrating that PRF-mediated autoregulation limits RF2 production in *P. fluorescens* SBW25.

We then assessed phenotypic consequences of removing *prfB* autoregulation. The *prfB*^ΔStop^ mutant grew similarly to wild type in different liquid media and formed colonies of equivalent sizes on plates (Figure 2C and Figure 3C). We also competed *prfB*^WT^ and *prfB*^ΔStop^ strains against a *prfB*^WT^ strain constitutively expressing *lacZ* (for colony discrimination on plates with X-gal) in co-culture and found no competitive disadvantage associated with removing the stop codon (Figure 2D). These results demonstrate that *P. fluorescens* carries a functional PRF system that mediates the autoregulation of RF2. Yet, elimination of the autoregulation has no detectable cost under the conditions tested.

**Figure 3.**
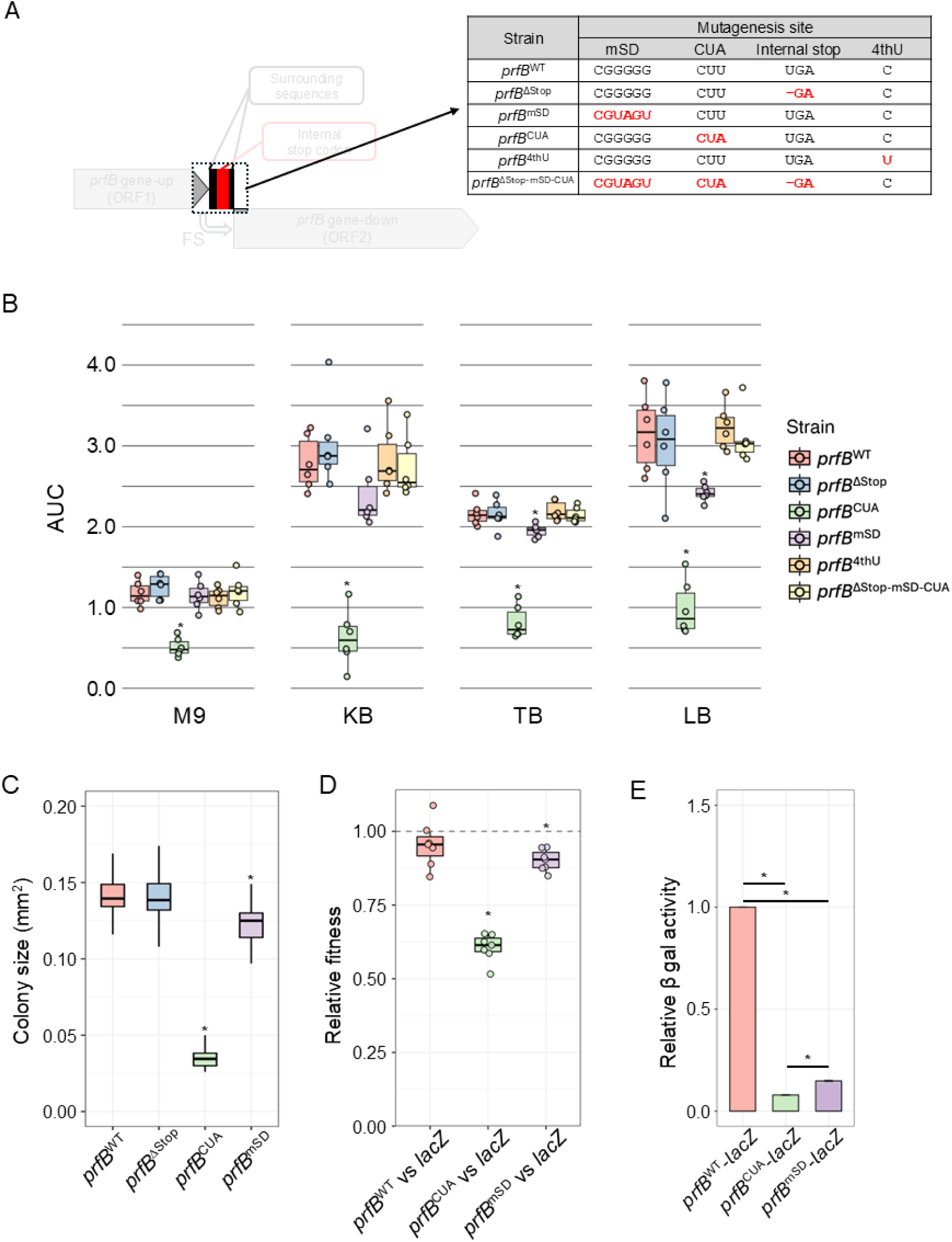
Characterization of mutations affecting conserved elements in the PRF site. (**A**) Diagram of mutations introduced at the PRF site. The table lists each allele with its nucleotide changes; mutated elements are shown in red. (**B**) Area under the growth curve (AUC) for *P. fluorescens* strains carrying the *prfB* alleles described in panel A. Cultures were grown in M9 minimal medium with 0.4% w/v Glucose (M9), King’s Broth (KB), Terrific Broth (TB), or Lysogeny Broth (LB). Box plots summarize six independent replicates (individual values shown as circles). See caption of Figure 2C for box plot description. Values significantly lower than *prfB*^WT^ (p ≤ 0.05) are indicated with an asterisk. (**C**) Colony size of strains carrying the indicated *prfB* alleles, after incubation on KB plates for 48 h at 28 °C. Colonies from three replicate plates were measured and averaged for each strain. Values significantly lower than *prfB*^WT^ (p ≤ 0.05) are indicated with an asterisk. (**D**) Competition assays of *P. fluorescens* strains carrying the wild-type *prfB* (*prfB*^WT^, orange), the *prfB*^CUA^ (green) or the *prfB*^mSD^ allele (purple) against an isogenic reference strain bearing wild-type *prfB* and constitutive *lacZ* (labelled as *lacZ*) for colony discrimination on X-gal plates. Assays were performed in King’s Broth (KB). A relative fitness of 1 is indicated by the dashed line. Box plots summarize seven independent replicates (individual values shown as circles). Values significantly lower than *prfB*^WT^ vs *lacZ* competition (p ≤ 0.05) are indicated with an asterisk. (**E**) β-galactosidase activity of *prfB*^WT^-*lacZ*, *prfB*^CUA^-*lacZ* and *prfB*^mSD^-*lacZ* fusions. Data were normalized to *prfB*^WT^-*lacZ*, and represent the average and standard error of five independent replicates. Statistically significant differences between value pairs are indicated with an asterisk.

### Fitness is impaired by mutations that reduce frameshifting rate

Next, we explored whether introduction of other modifications in the PRF site that might affect frameshifting rate caused noticeable phenotypes. We generated three additional variants (Figure 3A): (i) messy Shine-Dalgarno (*prfB*^mSD^), in which the last two G nucleotides of each triplet were replaced by U (from CGG GGG to CGU GGU), (ii) a change in the leucine codon within the shifty-stop sequence from CUU to the rarer leucine codon CUA (*prfB*^CUA^), which recruits a different tRNA-Leu (anticodon UAG rather than GAG (Chan and Lowe 2016)) at the P site and prevents wobble pairing with the +1 codon UUU, and (iii) a C-to-U substitution in the 4th position (*prfB*^4thU^). Importantly, none of these mutations alter the amino acid sequence of full-length RF2. Instead, they are predicted to affect termination efficiency at the internal stop codon and thereby modulate intracellular RF2 levels.

Two of the three modifications caused growth defects in liquid culture and produced smaller colonies on plates (Figure 3B and C). The defects were strongest in the *prfB*^CUA^ mutant and more modest in the *prfB*^mSD^ mutant, whereas the *prfB*^4thU^ mutant showed no detectable defect. We confirmed these trends by competitions with the wild type in co-culture (Figure 3D). We also observed that the strength of the growth defects was dependent on the medium (Figure 3B). In minimal medium (M9), the defects were milder (and even absent in the case of *prfB*^mSD^ mutant) than in rich media (KB, TB and LB).

We reasoned that the reduced growth of the *prfB*^CUA^ strain, and to a lesser extent *prfB*^mSD^ strain, reflected decreased RF2 production, as both mutations were predicted to lower frameshifting rates. Consistent with this, attempts to combine the CUA and mSD mutations yielded no transformants, indicating that the combination is highly deleterious, but the double mutant was readily obtained and grew like wild type in a background lacking the internal stop codon (Figure 3A-B). We confirmed reduced expression of RF2 using *lacZ* reporters and western blotting (Figure 3E and Supplementary Figure S1). In both mutants, RF2 protein levels dropped sharply and were barely detected by western blotting (Supplementary Figure S1). In *lacZ* reporter mutants, *prfB*^mSD^-*lacZ* and *prfB*^CUA^-*lacZ,* β-galactosidase was also strongly reduced, with the *prfB*^CUA^-*lacZ* showing a larger decrease, in agreement with its stronger growth defect (Figure 3E).

Overall, these results demonstrate that mutations that reduce frameshifting lower RF2 levels and reveal that the associated fitness costs are context-dependent, being more pronounced in rich medium than in minimal medium.

### Two types of mutations compensate the fitness defect caused by the *prfB*^CUA^ mutation

We next investigated whether secondary mutations could suppress the fitness defect caused by the *prfB*^CUA^ mutation. To this end, we propagated the mutant strain in KB medium with daily transfers (Figure 4A). Clones with increased fitness dominated the population after one to three weeks. We isolated a clone from each evolving population and sequenced their whole genomes to identify the compensatory mutations. To assess for parallelism in the outcomes, we repeated the experiment and sequenced candidate loci by Sanger sequencing (see Methods and Supplementary methods). In total, we characterized 34 isolates from independently evolved populations, which fell into two categories depending on whether the mutations were within the *prfB* PRF site (intragenic suppression, 4 isolates) or elsewhere in the chromosome (extragenic suppression, 30 isolates) (Table 1)

**Figure 4.**
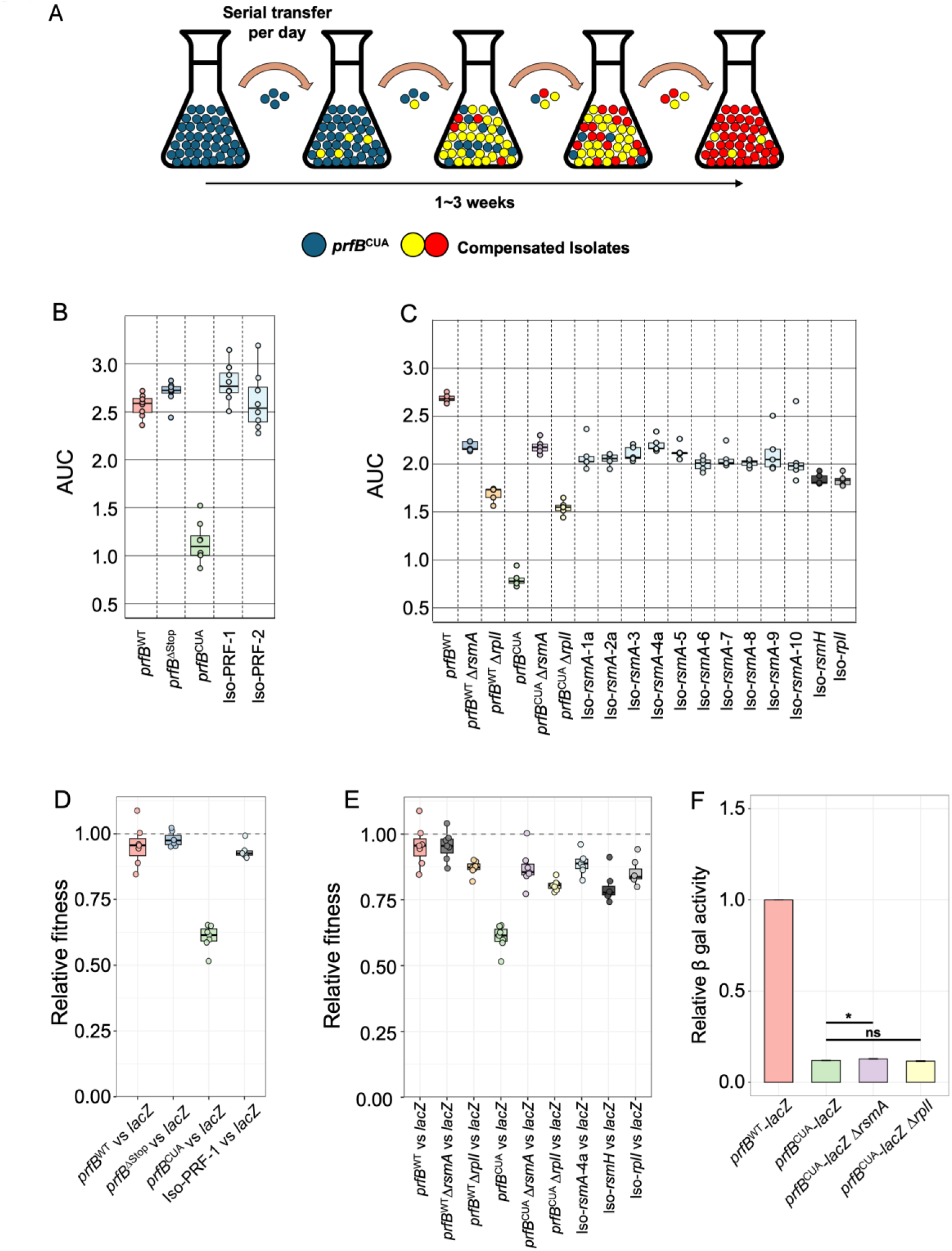
Isolation and characterization of mutations that compensate the *prfB*^CUA^ fitness defect. (**A**) Diagram of the serial-transfer evolution experiment. Populations were transferred every 24 h at 1:100 or 1:1000 dilution. Blue circles, ancestral *prfB*^CUA^ strain; yellow and red circles, the compensated mutants. Further details are in the Supplementary Methods (“Evolution Experiment List”). (**B**) Area under the growth curve (AUC) by 12 h for the following *P. fluorescens* strains: *prfB*^WT^ (orange)*, prfB*^Δstop^ (dark blue)*, prfB*^CUA^ (green), and *prfB*^CUA^ with intragenic suppressor alleles isolated from the evolution experiment (light blue) grown in KB medium. Data represents eight independent replicates. (**C**) AUC for *prfB*^WT^ or *prfB*^CUA^, alone or combined with Δ*rsmA*, Δ*rplI*, or extragenic suppressor alleles (light blue), growing in KB medium. Data represents five independent replicates. (**D**) Competition assays of *prfB*^WT^ (orange), *prfB*^ΔStop^ (dark blue), *prfB*^CUA^ (green) or *prfB*^CUA^ with a representative intragenic suppressor (Iso-PRF-1, light blue) against an isogenic *prfB*^WT^ reference strain bearing constitutive *lacZ* (labelled as *lacZ*) for colony discrimination on X-gal plates. Assays were performed in KB medium. A relative fitness of 1 is indicated by the dashed line. Data represents eight independent replicates. (**E**) Competition assays of *prfB*^WT^ or *prfB*^CUA^, alone or with Δ*rsmA*, Δ*rplI*, or representative extragenic suppressor alleles, against the same reference. Data represents five independent replicates. (**F**) β-galactosidase activity of *prfB*^WT^-*lacZ* and of *prfB*^CUA^-*lacZ* fusions in wild-type, Δ*rsmA* or Δ*rplI* backgrounds. Data were normalized to *prfB*^WT^-*lacZ*, and represent the average and standard error of five replicates. Statistically significant differences between value pairs are indicated with an asterisk.

**Table 1.**
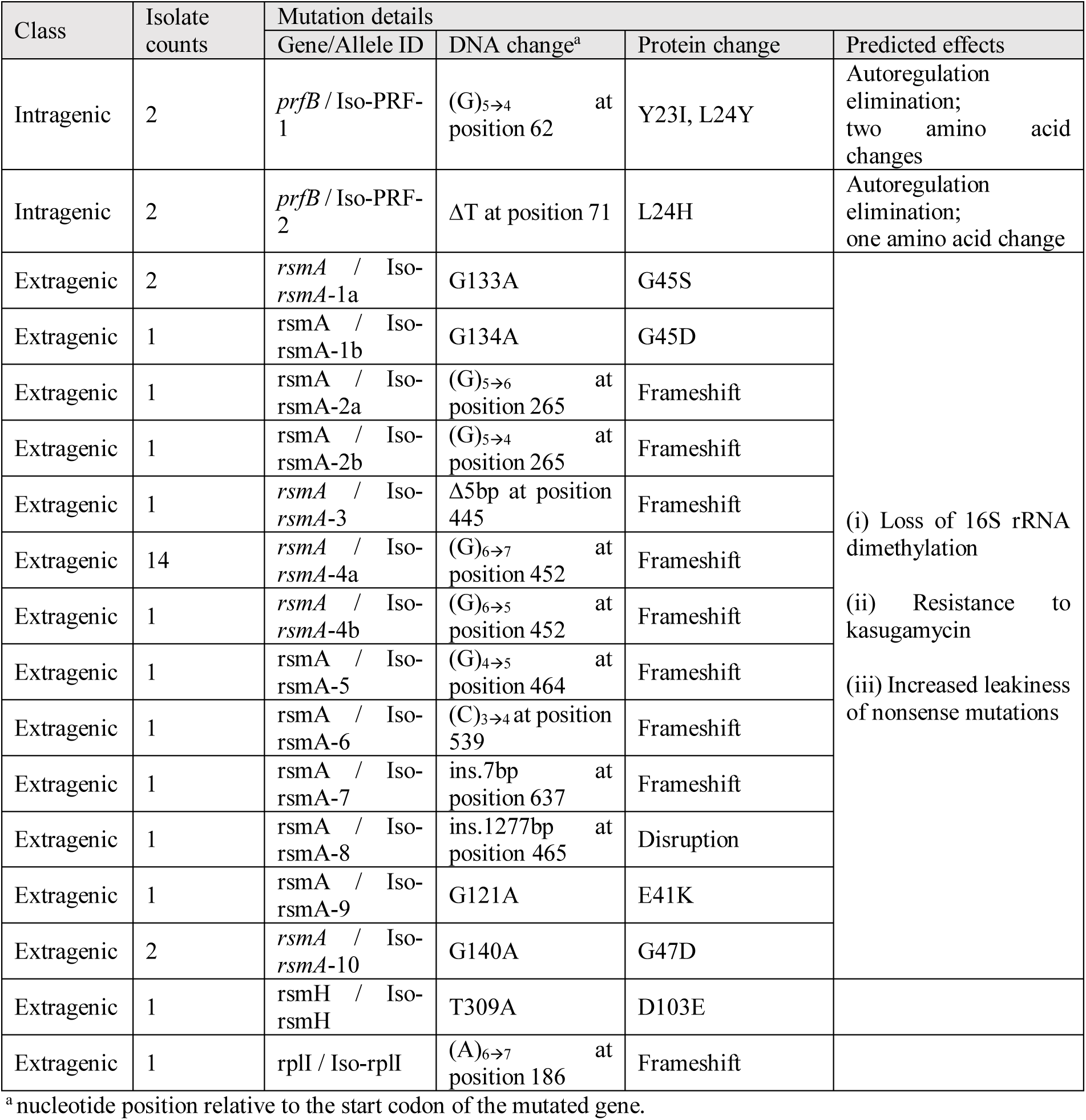
Mutations increasing fitness of *prfB*^CUA^ isolated through experimental evolution.

The intragenic suppressors harbored single nucleotide deletions at the PRF site. Two isolates had a deletion of a G in the SD-like sequence, and two others a deletion of a T (U) immediately preceding the internal stop codon. Both types of mutations bypassed the internal stop codon and enabled RF2 production without the need for frameshifting. This compensation effectively abolished *prfB* autoregulation, resulting in elevated RF2 levels comparable to those of the *prfB*^ΔStop^ mutant (Supplementary Figure S1), and restored growth and fitness to wild-type levels (Figure 4B and D).

Most suppressors (30/34) were extragenic. The majority carried mutations in *rsmA* (also known as *ksgA*) (Table 1), a gene encoding a highly conserved 16S rRNA methyltransferase required for proper ribosome assembly (Pletnev *et al*. 2020). Loss of RsmA has been shown to confer resistance to the aminoglycoside antibiotic kasugamycin (Helser, Davies and Dahlberg 1972; Van Buul, Visser and Van Knippenberg 1984). These mutations included missense substitutions as well as multiple insertions and deletions that altered the reading frame (Table 1). Notably, 14 isolates carried a single-nucleotide expansion in a polyG tract in the middle of the gene, consistent with impaired RsmA function (Table 1). Accordingly, representative isolates were resistant to kasugamycin (Supplementary Figure S2). In addition, one clone carried a single point mutation in *rsmH*, another 16S rRNA methyltransferase, resulting in a D103E amino acid substitution, and one clone carried a frameshift mutation in *rplI*, which encodes the ribosomal protein L9 (Table 1).

To evaluate the compensation efficiency of these mutations, we monitored growth of representative *rsmA*, *rsmH* and *rplI* isolates and performed competition assays against a wild type ancestral constitutively expressing lacZ for colony discrimination on X-gal (Figure 4C and E). As controls, we included *rsmA* and *rplI* deletion mutants (we were unable to obtain a *rsmH* deletion mutant). All mutations improved the growth and fitness of the *prfB*^CUA^ strain, with *rsmA* mutations providing the strongest rescue. However, both *rsmA* and *rplI* deletion decreased growth compared to the wild type, with *rplI* defect being more severe.

The inactivation of RsmA, RsmH or RplI has previously been reported to reduce translational fidelity and promote more frequent frameshifting on UGA stop codons (Van Buul, Visser and Van Knippenberg 1984; Herbst *et al*. 1994; Herr *et al*. 2001; Kimura and Suzuki 2010; Smith *et al*. 2019). We therefore hypothesized that these compensatory mutations restored growth of the *prfB*^CUA^ by increasing frameshifting rate at the PRF site of *prfB*, thereby increasing RF2 levels. To test this, we first measured RF2 levels by western blot in the *prfB*^CUA^ background with and without *rsmA* or *rplI* deletions. In all cases, RF2 levels remained low, barely above detection limit (Supplementary Figure S1), suggesting that inactivation of RsmA or RplI did not substantially increase frameshifting rates to wild type levels. We further explored this by measuring β-galactosidase activity using *prfB*^CUA^-*lacZ* fusions in wild type, RsmA^−^ or RplI^−^backgrounds (Figure 4D). Only a mild (but statistically significant) increase of β-galactosidase activity was observed in the *rsmA* mutant, and no increase in *rplI* mutant. Together, these results indicate that the rescue of the *prfB*^CUA^ strain by mutations in *rsmA* and *rplI* is not exclusively due to increased frameshifting at the PRF site of the *prfB* gene.

### Phylogenetic analysis reveals no significant correlation between genomic context and presence of PRF site in *prfB*

The observation that mutations abolishing *prfB* autoregulation are selected in mutants with reduced frameshifting rates (Figure 4) is compatible with the idea that evolutionary loss of *prfB* autoregulation is favored when RF2 becomes limiting. In this context, previous proposals for the maintenance and loss of *prfB* autoregulation across bacterial species have emphasized correlations with genomic features. For example, high usage of the UGA stop codon has been proposed to favor autoregulation loss by increasing the demand for RF2 (Prince, Lin and Feaga 2025). Conversely, maintenance of autoregulation may be favored by features that increase the cost of RF2 overactivity. For instance, RF2 alleles with enhanced ability to recognize UAG stop codons have also been shown to misrecognize UGG tryptophan codons in *S. enterica*, increasing premature termination at these codons (Abdalaal *et al*. 2020). Thus, high UGG codon usage could also contribute to the maintenance of *prfB* autoregulation in some cases.

To explore these possibilities, we analyzed the genomes of 818 bacterial species from 35 phyla, including both species that retain (PRF^+^) and that have lost (PRF^−^) *prfB* autoregulation (Supplementary Table S2 and S3) (sequences kindly provided by Dr. Fredrick (Naeem *et al*. 2023)). We reconstructed the phylogeny based on 81 core gene sequences (Kim *et al*. 2021), and mapped the PRF status of each species onto the tree (Figure 5A). This analysis indicated that autoregulation loss occurred multiple times independently across bacterial phyla, consistent with previous reports (Naeem *et al*. 2023; Prince, Lin and Feaga 2025). However, autoregulation loss was not homogeneously distributed across the tree, but instead clustered within clades (Figure 5A), suggesting that once lost, it was largely vertically inherited and that regain events were rare.

**Figure 5.**
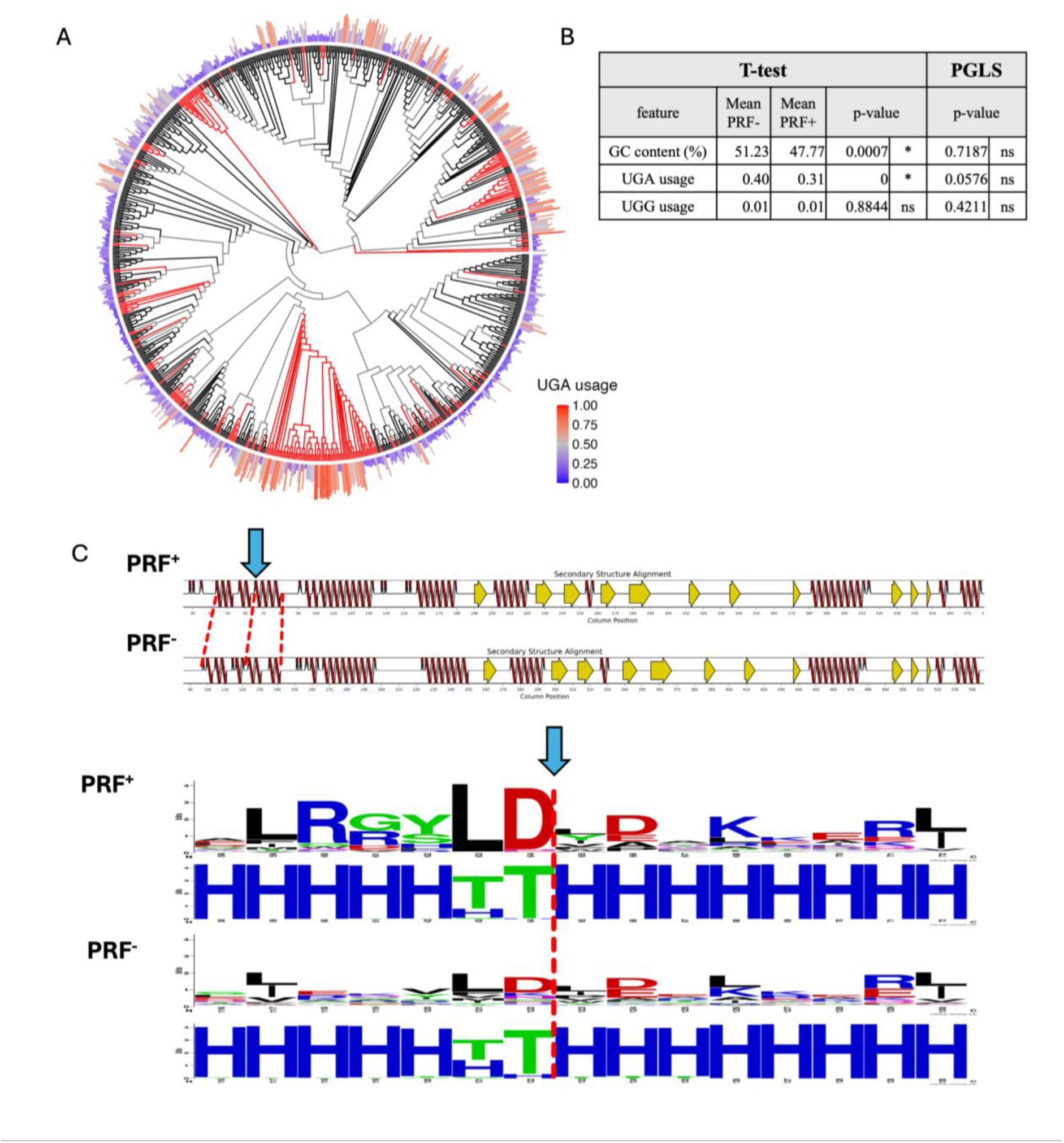
Computational analysis of PRF sites across bacterial species. **(A)** Phylogenetic tree of 818 bacterial species. Leaves and branches for PRF⁺ species are black; PRF⁻ species are red. Species-level UGA stop codon usage is represented as a color-coded histogram, with a frequency of 0.5 shown in gray, higher frequencies in red, and lower frequencies in blue. The tree was built using UBCG2 (Kim et al. 2021). See Supplementary Figure S4 for a high-resolution version of the tree including species names. **(B)** Differences in GC content, UGA stop codon usage and UGG tryptophan codon usage between PRF^+^ and PRF^−^ species. For each feature, differences between PRF⁺ and PRF⁻ groups were tested with Welch’s t-test and with Phylogenetic Generalized Least Squares (PGLS), a method that accounts for non-independence among species. Significance: * p ≤ 0.05; ns p > 0.05. p-values < 1×10⁻⁴ are displayed as 0. **(C)** Predicted consensus secondary structure of the PRF region within RF2 in PRF⁺ species (n = 575) and of the equivalent region in PRF⁻ species (n = 243) (SIMSApiper v2). Red lines, α-helices; yellow arrows, β-strands; black lines, loops or alignment gaps. The blue arrow marks the PRF site in PRF⁺ species. Red dashed lines connect the start and end of the first two α-helices in each group. Bottom, WebLogo plots derived from the corresponding amino-acid alignments (top for each group) and DSSP-based secondary-structure strings (bottom for each group); the blue arrow (PRF⁺) and red dashed line (PRF⁻) indicate the PRF site/orthologous region.

We then tested if PRF status correlates with genomic features that could influence the maintenance or loss of *prfB* autoregulation (Supplementary Table S4). Specifically, we examined usage of the UGG tryptophan codon or the UGA stop codon. UGG usage was almost identical in PRF^+^ and PRF^−^ species, suggesting that it does not explain the retention or loss of autoregulation (Figure 5B). However, PRF^+^ and PRF^−^ species differed in UGA usage: UGA accounted for 31% of stop codons in PRF^+^ species and ∼40% in PRF^−^ species, consistent with earlier observations (Figure 5B) (Prince, Lin and Feaga 2025). While this pattern could suggest that increased UGA usage favors autoregulation loss, an alternative explanation is that it reflects phylogenetic clustering, as both PRF status and UGA usage tend to be conserved within clades (Figure 5A). To account for phylogenetic relatedness, we applied phylogenetic generalized least squares (PGLS) analysis. This analysis reduced the strength of the association between UGA usage and PRF status and did not reach statistical significance at conventional thresholds (p = 0.0576) (Figure 5B), indicating that the apparent correlation is largely explained by shared ancestry, while any remaining effect is at most weak. A close inspection of the tree supported this interpretation: while some autoregulation loss events were associated with higher UGA codon usage than neighboring PRF^+^ species (see, for example, *Faecalibaculum rodentium* in the high-resolution tree with species names provided in Supplementary Figure S3), this association was not universal, and we observed many cases of high-UGA species that retain autoregulation and, conversely, species with low UGA usage that have lost autoregulation.

## DISCUSSION

Here, we have characterized PRF in the *prfB* gene of *P. fluorescens* and investigated the evolutionary dynamics of mutants with reduced frameshifting. Our data indicate that *prfB* autoregulation is dispensable under laboratory conditions, but that mutations that reduce frameshifting rate impose a context-dependent fitness cost, which is stronger in rich media than in minimal media. Using experimental evolution, we have found that bacteria can escape this constraint either by inactivating accessory ribosomal proteins or by a single-nucleotide deletion immediately upstream of the internal stop codon in *prfB*, which bypasses the need for PRF to produce full-length RF2.

These results suggest that increasing RF2 demand (or reduced RF2 availability) can act as a trigger for *prfB* autoregulation loss. We envision three non-mutually-exclusive scenarios where this could happen. First, higher RF2 demand may be imposed by the genomic context. One obvious possibility, in line with previous proposals (Wei, Wang and Xia 2016; Ho and Hurst 2022; Prince, Lin and Feaga 2025), is that elevated usage of the UGA codon increases the cellular requirement for RF2 and thereby favors mutants in which autoregulation is eliminated. Under this view, lineages with greater reliance on UGA would experience more frequent or more severe bouts of RF2 limitation, making mutations that eliminate *prfB* autoregulation advantageous. A straightforward prediction is a positive association between autoregulation loss and UGA usage. Indeed, comparisons of PRF^+^ and PRF^−^ species show higher mean UGA usage in PRF^−^ lineages (Figure 5B), consistent with earlier observations (Prince, Lin and Feaga 2025). However, when we accounted for shared ancestry using PGLS, this association was not significant (Figure 5B), suggesting that elevated UGA usage alone does not impose a sufficiently strong or consistent RF2 demand to leave a robust phylogenetic signal in our dataset. Moreover, several PRF^−^ clades exhibit low UGA usage (Figure 5A; Supplementary Figure S3), indicating that additional evolutionary pressures must be at play. It remains possible, however, that the relevant signal is transcriptomic or locus-specific rather than genomic: for example, enrichment of UGA stops within a subset of highly expressed genes could create a substantial RF2 burden without shifting genome-wide averages, or the presence of an inefficient UGA in a particular gene could, when mistranslated, produce a toxic protein variant that perturbs physiology and intensifies selection for autoregulation loss. Such localized or transcriptomic pressures would be difficult to detect with global codon-usage summaries and may leave little phylogenetic trace, yet they could still provide the immediate genetic conditions that tip lineages toward the loss of *prfB* autoregulation.

A second scenario that could impose a higher requirement for RF2—and thereby favor autoregulation loss—is ecological. Exposure to particular antibiotics or toxic molecules can elevate termination demand. Aminoglycosides, for example, are known to increase stop codon readthrough (Diop, Chauvin and Jean-Jean 2006; Malik *et al*. 2010), a defect that could be ameliorated by higher levels of translation termination factors. In this sense, it has been recently reported that a *Flavobacterium johnsoniae* mutant lacking *prfB* autoregulation exhibits a modest growth advantage when exposed to sublethal concentrations of streptomycin (Naeem *et al*. 2023), suggesting that increased RF2 levels can contribute to aminoglycoside resistance in at least some contexts. Consistent with this, we observed that the lack of *prfB* autoregulation causes a modest increase in resistance to the aminoglycosides streptomycin and gentamicin in *P. fluorescens* SBW25 (Supplementary Figure S4). In contrast, no significant change was observed for chloramphenicol, which inhibits peptide elongation thus stalling translation elongation, indicating that this phenotype is aminoglycoside-specific (Supplementary Figure S4). More broadly, ecological regimes that entail sustained elevations in global gene expression, such as frequent nutrient upshifts or oscillatory feast-famine cycles, could also select for higher RF2 levels by increasing overall translation flux, thereby creating periods of RF2 limitation and, in turn, favoring the loss of autoregulation.

The third scenario that may favor autoregulation loss is analogous to our evolution experiment: the gradual accumulation of mutations in the *prfB* PRF element that lower frameshifting efficiency, whose fitness costs can be suppressed by eliminating autoregulation altogether. Such mutations could be nearly neutral when the translational burden is low, for example during slow growth. Indeed, our results show that the growth defects caused by the *prfB*^CUA^ and *prfB*^mSD^ alleles are smaller in minimal than in rich medium, with *prfB*^mSD^ showing no defect in minimal medium (Figure 3B), suggesting that the fitness cost associated with lower RF2 levels is context dependent. However, if conditions change and translation rate increases, low-frameshifting mutants may not be able to cope with the increased translation termination demand, leading to the selection of suppressors that restore fitness. As shown in our experiments, some of these suppressors are single-nucleotide deletions upstream of the internal stop codon in *prfB*, which bypass the internal stop codon and eliminate autoregulation.

Any of the three scenarios outlined above could increase RF2 demand, thereby favoring autoregulation loss. However, PRF itself may be under direct positive selection in some species or ecological contexts, as it enables cells to tune RF2 abundance to match translational demands within a certain range. In this sense, overproduction of RF2 has been shown to be detrimental in some species under certain laboratory conditions (Jørgensen *et al*. 1993; Rengby and Arnér 2007; Abdalaal *et al*. 2020; Lalanne, Parker and Li 2021; Prince, Lin and Feaga 2025), and the same is likely true in natural environments. Thus, autoregulation loss via the compensatory routes we describe should be selectively favored only when increased RF2 availability is strongly beneficial (for example, under RF2-limiting conditions) and the fitness costs of RF2 overproduction and loss of autoregulation are comparatively small.

In PRF^‒^ species, the nucleotide and amino acid sequences surrounding the PRF site (from the Shine–Dalgarno-like sequence to the internal stop codon) are degraded (Prince, Lin and Feaga 2025) (Figure 5C). This neutral sequence degradation likely hinders the regain of autoregulation, because it is expected to disrupt elements required to sustain high frameshifting rates. Thus, regain events, if any, would most likely occur shortly after loss, before the surrounding PRF region decays, or via horizontal gene transfer from a PRF^+^ donor. Despite the decay of the amino acid sequence, the predicted secondary structure—consisting of two initial α-helices and the intervening loop—is retained, suggesting that this structural element is under selection independently of PRF status. This observation is consistent with the proposal that the PRF site is positioned to minimize its additional structural impact on RF2 translation (Worth, Gong and Blundell 2009).

In addition to eliminating *prfB* autoregulation, our evolution experiment repeatedly produced mutations that inactivated or reduced the function of the accessory ribosomal proteins RsmA, RsmH, and RplI (Table 1). Although loss of these proteins is not an optimal solution in our experimental conditions, as evidenced by the reduced fitness of the Δ*rsmA* and Δ*rplI* mutants relative to wild type (Figure 4C), such mutations arose in many independent lineages, suggesting they are highly accessible. Mutations in *rsmA* were especially frequent. In particular, indels in poly-G tracts within the coding sequence caused frameshifts and complete loss of RsmA function (Table 1). This pattern raises the possibility that *rsmA* is a mutational hotspot. Such a bias might have an evolutionary explanation: RsmA deficiency confers resistance to the antibiotic kasugamycin (Helser, Davies and Dahlberg 1972; Van Buul, Visser and Van Knippenberg 1984) (Supplementary Figure S2) and can ameliorate genetic perturbations that impair translation initiation (Das *et al*. 2008). Since poly-G tracts are prone to slippage (Levinson and Gutman 1987; Burch, Danaher and Stein 1997), the ability to expand or contract these tracts could create locally elevated, reversible mutability, allowing transient inactivation of *rsmA* during periods of translational stress and restoration of function once the perturbation subsides.

The mechanisms underlying suppression of the *prfB*^CUA^ growth defect by mutations in *rsmA*, *rsmH* and *rplI* remain unclear. Inactivation of RsmA or RplI produced only minor or negligible increases in *prfB* frameshifting (Figure 4F; Supplementary Figure S1), indicating that suppression is unlikely to result from restoration of sufficient frameshifting rate and consequent elevation of RF2. An alternative, non-exclusive explanation is that loss of these accessory factors increases the probability of bypassing UGA stops (i.e., enhanced readthrough or miscoding at termination), which has been reported for RsmA^‒^ mutants in some contexts (Van Buul, Visser and Van Knippenberg 1984). Under low-RF2 conditions, ribosomes are expected to dwell abnormally long at UGA codons, creating queues that sequester ribosomes and limit recycling for new rounds of initiation. If inactivation of RsmA or RplI raises the rate of UGA bypass (or otherwise reduces dwell time at problematic stops) this would lessen ribosome queuing, increase the pool of recyclable ribosomes, and partially restore growth despite persistent RF2 scarcity. By relieving queuing, these changes may also mitigate secondary consequences of RF2 depletion on global mRNA stability. Specifically, because translating ribosomes protect bacterial mRNAs from degradation (Régnier and Arraiano 2000; Müller *et al*. 2025; Zhang *et al*. 2025), ribosomes trapped at UGA codons on a subset of transcripts would otherwise deplete ribosome density elsewhere, rendering many mRNAs more prone to decay. A second, complementary possibility is a general effect on translation capacity: RsmA loss perturbs 30S maturation and initiation fidelity (Sharma and Anand 2019), and RplI influences elongation dynamics and quality control (Herbst *et al*. 1994). The inactivation of either protein could reduce effective initiation throughput or alter elongation kinetics enough to lower the overall demand on termination, thereby rebalancing translational demand and RF2 availability. Together, these effects would relieve the bottleneck created by RF2 limitation without directly restoring frameshifting. Further work is necessary to explore these mechanisms.

Overall, our results clarify the mechanisms of RF2 autoregulation in *P. fluorescens* and show how changes in RF2 demand, coupled with compensatory mutations in *prfB*, can drive the loss of autoregulation. Altogether, this work helps explain the maintenance and repeated loss of *prfB* autoregulation across bacterial lineages.

## METHODS

### Strains and growth conditions

All the strains used in this study are derivatives of *P. fluorescens* SBW25 (NCBI Reference Sequence: NC_012660.1) (Silby *et al*. 2009). Strain and oligonucleotide lists are provided as Supplementary Table S1. All mutants were constructed using scar-free two-step allelic exchange protocol using pUI*sacB*, as previously described (Farr *et al*. 2025). King’s medium B (KB) (King, Ward and Raney 1954), Lysogeny broth (LB), Terrific broth (TB) or M9 minimal media with 0.4% w/v Glucose were used for culturing bacteria, depending on the experiment. Strains were routinely grown on KB agar plates at 28°C for 48 h. Liquid cultures were grown overnight at 28°C in an orbital shaking incubator at 220 rpm.

### β-galactosidase assays

β-galactosidase assays were performed as previously described (Raval *et al*. 2023), with modifications for strains with low *lacZ* expression. To increase sensitivity, we extended the reaction time up to 24 h and concentrated overnight cultures fivefold. Z buffer (Na_2_HPO_4_ 60 mM, NaH_2_PO_4_ 40 mM, KCl 10 mM, MgSO_4_ 1 mM, 5% β-mercaptoethanol) was freshly prepared before each assay. Chloramphenicol (100 µl, 3 mg/ml) was added to 500 µl of concentrated overnight culture to block translation. After 10 min on ice, 350 µl of Z buffer was added and incubation on ice was continued for 1 h. The reaction was initiated by adding 200 µl of o-nitrophenyl-β-D-galactopyranoside (ONPG) solution (12 mg/ml) and incubating at 30°C. At each time point, 50 µl of the reaction mixture was withdrawn and the reaction was stopped by adding 25 µl of 1 M Na_2_CO_3_. After centrifugation (12,000 g, 1 min), 50 µl of the supernatant was transferred to a clear 96-well plate and OD_420_, OD_550_ and OD_600_ were measured. Because of the long incubation times, β-galactosidase activity was compared directly using OD_420_ values after confirming the absence of cell debris in the supernatant by OD_550_ and OD_600_ measurements.

### Growth curves and competition assays

For growth curves, overnight precultures were prepared in 96-well plates by inoculating individual colonies into 200 µl of growth medium per well. Then, 2 µl of each preculture were inoculated into 198 µl of fresh growth medium in a 96-well plate. OD_600_ was recorded every 10 min, with orbital shaking (282 rpm, 3 mm amplitude) for 5 s before each measurement.

Competition assays were performed as described previously (Khomarbaghi *et al*. 2024), in four experimental blocks with three replicate competitions per strain pair. Precultures were prepared as above. After overnight growth, the two competitors for each pair were mixed at an approximate 1:1 ratio, and 4 µl of this mixture were used to inoculate 4 ml KB in a 13-ml tube, which was then incubated for 24 h (28 °C, 220 rpm). Samples were taken at the start (T0) and end (T24) of the competition, serially diluted, and plated on LB agar containing 60 µg ml^−1^ X-gal (48 h, 28 °C). Colonies of each competitor were counted at T0 and T24, with strains distinguished by colony colour as one of the competitors was the SBW25-*lacZ* strain (Zhang and Rainey 2007), which expresses *lacZ* constitutively. Changes in competitor ratios were used to calculate relative fitness as described previously (Khomarbaghi *et al*. 2024).

### Colony size measurement

To measure the colony size of wild type and mutant strains, dilutions from overnight cultures were spread onto KB agar plates to obtain between 10 and 50 colonies per plate. The plates were incubated for 48 h at 28°C. After incubation, images of the plates were captured, and colony sizes were quantified using ImageJ (version 1.54p). For quantification, images were converted to 8-bit grayscale and segmented using a fixed pixel intensity threshold of 75–255. We filtered out the segmentations below 0.015 mm^2^ and applied circularity filter of 0.80–1.00 to exclude non-colony artifacts.

### Serial transfer evolution experiment

Individual colonies of the *prfB*^CUA^ strain were used to inoculate 4 ml KB medium in 13-ml tubes to generate precultures. For each transfer, 40 µl or 4 µl of preculture were inoculated into 4 ml fresh KB medium in a new 13-ml tube, and each lineage was subsequently transferred with the same volume into fresh medium every 24 h. Colony Forming Units (CFU) assays were performed on day 1 and then weekly to monitor contamination and the appearance of compensatory mutations (assessed by colony size). Cryo-stocks of the evolving populations were prepared every week and stored at −80 °C. From the weekly CFU plates, a single colony per lineage was picked after 48 h incubation at 28 °C. When colonies of different sizes were present, we selected the largest colony; otherwise, a colony of median size was chosen. As a control for contamination, a fresh tube containing only KB medium was carried through the daily transfer regime.

### Sequencing

Evolved isolates showing significant growth improvement were first Sanger-sequenced across the *prfB* coding region. Isolates without mutations in *prfB* were then subjected to whole genome sequencing (detailed in the next paragraph). After *rsmA* was identified as a recurrent mutational target in the evolution experiment, both *prfB* and *rsmA* were Sanger-sequenced in all selected evolved isolates. Whole-genome sequencing was additionally performed for isolates that did not carry mutations in both gene. Primers used for Sanger sequencing are listed in Supplementary Table 1. Geneious Prime (v 2025.1.3) was used to identify mutation sites.

Six isolates obtained from the evolution experiment were whole-genome sequenced (rsmA-1a-1, rsmA-2b, rsmA-4a-1, rsmA-5, rsmH-1, rplI-1). Library preparation and sequencing were performed by Novogene Europe (Cambridge, UK) using Illumina technology on a NovaSeq 6000 platform. For four isolates (rsmA-1a-1, rsmA-2b, rsmA-4a-1, rsmA-5), 150-bp paired-end reads were generated. For the remaining two isolates (rsmH-1, rplI-1), 250-bp paired-end reads were generated with an Illumina MiSeq instrument at the Max Planck Institute for Evolutionary Biology (Plön, Germany) using standard procedures (Khomarbaghi *et al*. 2024). A minimum of 0.575 million raw reads per genotype were obtained and aligned to the SBW25 reference genome (NC_012660.1) (Silby *et al*. 2009) using breseq (Deatherage and Barrick 2014) and Geneious Prime (v 2024.0.2).

### Western blotting

For each strain, a single colony from a 48-h plate was used to inoculate culture in 4ml of KB medium and grown overnight at 28°C in an orbital shaking incubator at 220 rpm. One ml of overnight culture was adjusted to OD_600_ = 2.0, centrifuged (12,000 g, 1 min), and the pellet resuspended in 200 µl sample preparation buffer (100 µl 2× Laemmli buffer, Bio-Rad #1610737; 95 µl distilled water; 5 µl β-mercaptoethanol). Samples were incubated at 95 °C for 5 min and centrifuged again (12,000 g, 10 min).

Stacking (4% polyacrylamide) and running (10% polyacrylamide) gels were cast following the Handcasting Polyacrylamide Gels protocol (Bio-Rad Bulletin 6201) and mounted in a Mini-PROTEAN Tetra Vertical Electrophoresis Cell (Bio-Rad #1658004). Fifteen µl of each sample or protein ladder (Bio-Rad #1610376) were loaded per lane. Electrophoresis was carried out at 30 V for 30 min followed by 70 V for 150 min. Proteins were then transferred to a PVDF membrane (Bio-Rad #1704156) using the Trans-Blot Turbo Transfer System (Bio-Rad #1704150).

Membranes were blocked for 1 h at room temperature in 5% skim milk in Tris-buffered saline with 0.1% Tween 20 (TBST), then incubated overnight at 4 °C with primary antibody diluted to a final concentration of 0.5 µg/ml in 5% skim milk in TBST. The primary antibody was a rabbit polyclonal antibody raised against a *P. fluorescens* SBW25 RF2 antigenic peptide (CVRKSPFDSGNRRHT) by GenScript. After five 10-min washes in TBST, membranes were incubated for 1 h at room temperature with secondary antibody (goat anti-rabbit IgG–HRP, Bio-Rad #1721019) diluted 1:3000 in 5% skim milk in TBST, followed by five additional 10-min washes in TBST. All incubation and rinsing steps were performed with gentle agitation at ∼30 rpm. Chemiluminescent substrate (Abcam #ab133406) was applied for 2 min before imaging, and signals were captured using a Bio-Rad ChemiDoc Imaging System.

### Phylogenetic analyses

RF2 amino acid sequence and its PRF status was kindly provided by Dr. Fredrick’s group at Ohio State University (Naeem *et al*. 2023). A database of 4971 microbial genomes was downloaded from NCBI genome dataset filtered with (i) reference genomes, (ii) annotated genomes, (iii) assembly level: Complete at 04.2023. Detailed species list is provided in Supplementary Table S2. The genome database was dereplicated using dRep v.3.5.0 to reduce redundancy (over-representation of closely related genomes) and to filter for high-quality genome assemblies (Olm *et al*. 2017). Filtering was performed with the following criteria: Completeness ≥ 90% and contamination ≤ 1% (ran by CheckM taxonomy workflow); primary clustering threshold of 70% ANI; secondary clustering threshold of 50% ANI using the fastANI algorithm. The list of genomes retained at each filtering step, together with the full filtering workflow, is provided in Supplementary Table S3.

Phylogeny was reconstructed based on 81 core genes, using the UBCG2 pipeline (Kim *et al*. 2021). Weblogo image showing amino acids and secondary structures at the PRF site was constructed based on an alignment of RF2 amino acid sequences generated with SIMSApiper v2, using Weblogo webtool (https://weblogo.berkeley.edu/logo.cgi).

Phylogenetic generalized least squares (PGLS) analysis was performed to test whether PRF status is associated with genomic features such as codon usage and GC content. PRF status was used as the predictor variable, coded as 0 for PRF− and 1 for PRF+. The following response variables were tested: (i) GC content (percentage, 0–100), (ii) UGA stop codon usage (relative frequency among the three stop codons, 0–1), and (iii) UGG codon usage (frequency relative to total sense codons, 0–1). Feature values for each species are provided in Supplementary Table S4. PGLS was performed using the phylolm package under a Brownian motion (BM) evolutionary model. For non-phylogenetic comparisons, Welch’s two-sample t-tests (unequal variances) were used. Statistical significance was defined as p < 0.05. Version information for the software and packages used for phylogenetic analyses were as follows: R version: 4.4.1 (2024-06-14); R package("phylolm"): 2.6.5; R package("ape"): 5.8.1; R package ("ggplot2"): 3.5.1

### Statistical analyses

All statistical analyses were performed in R version 4.4.1 (2024-06-14), using the “stats” package. Unless otherwise stated, differences between groups were assessed with Welch’s t-test. Normal p-values were reported for single pairwise comparisons while the Benjamini-Hochberg adjustment was applied for multiple comparisons. For the competition assays, we used a one-way ANOVA to test for differences among competition groups. Residuals violated normality (Shapiro–Wilk W = 0.952, p = 0.00337) while variances were homogeneous (Levene’s test F = 0.82, p = 0.62). Pairwise comparisons were then performed using Dunn’s test with Bonferroni correction to obtain adjusted p-values.

## ACKNOWLEDGEMENTS

The authors thank Dr. Kurt Fredrick for providing species list with ARFA analysis (Naeem *et al*. 2023), Sophie-Luise Heidig for providing useful information on performing SIMSApiper v2, Julien Dutheil for advising on phylogenetic analyses and Jessica Voꞵ for supporting evolution experiments.

## Author contributions

Conceived and designed the experiments: S.L., F.B., J.G. Performed the experiments: S.L. Analyzed the data: S.L., F.B., J.L.G., J.G. Wrote the paper: S.L., J.L.G., J.G.

## DATA AVAILABILITY

All raw experimental data including sequencing data can be found in Zenodo (doi:10.5281/zenodo.18065983).

## SUPPLEMENTARY MATERIALS

Supplementary Figures S1-S4 and Supplementary methods are provided in Supplementary.pdf, Supplementary tables Table S1-S4 are provided in Supplementary.xlsx

## REFERENCES

Abdalaal H, Pundir S, Ge X et al. Collateral toxicity limits the evolution of bacterial release factor 2 toward total omnipotence. Mol Biol Evol 2020;37:2918–30.

Adamski FM, Donly BC, Tate WP. Competition between frameshifting, termination and suppression at the frameshift site in the *Escherichia coli* release factor-2 mRNA. Nucl Acids Res 1993;21:5074–8.

Antonov I, Coakley A, Atkins JF et al. Identification of the nature of reading frame transitions observed in prokaryotic genomes. Nucl Acids Res 2013;41:6514–30.

Baranov PV, Gesteland RF, Atkins JF. Release factor 2 frameshifting sites in different bacteria. EMBO Reports 2002a;3:373–7.

Baranov PV, Gesteland RF, Atkins JF. Recoding: translational bifurcations in gene expression. Gene 2002b;286:187–201.

Baranov PV, Gesteland RF, Atkins JF. P-site tRNA is a crucial initiator of ribosomal frameshifting. RNA 2004;10:221–30.

Bekaert M, Atkins JF, Baranov PV. ARFA: a program for annotating bacterial release factor genes, including prediction of programmed ribosomal frameshifting. Bioinformatics 2006;22:2463–5.

Betney R, De Silva E, Krishnan J et al. Autoregulatory systems controlling translation factor expression: Thermostat-like control of translational accuracy. RNA 2010;16:655–63.

Biziaev N, Sokolova E, Yanvarev DV et al. Recognition of 3′ nucleotide context and stop codon readthrough are determined during mRNA translation elongation. J Biol Chem 2022;298:102133.

Brown CM, Dalphin ME, Stockwell PA et al. The translational termination signal database. Nucl Acids Res 1993;21:3119–23.

Brown CM, Stockwell PA, Dalphin ME et al. The translational termination signal database (TransTerm) now also includes initiation contexts. Nucl Acids Res 1994;22:3620–4.

Burch CL, Danaher RJ, Stein DC. Antigenic variation in Neisseria gonorrhoeae: production of multiple lipooligosaccharides. J Bacteriol 1997;179:982–6.

Caliskan N, Wohlgemuth I, Korniy N et al. Conditional Switch between Frameshifting Regimes upon Translation of dnaX mRNA. Molecular Cell 2017;66:558–567.e4.

Chan PP, Lowe TM. GtRNAdb 2.0: An expanded database of transfer RNA genes identified in complete and draft genomes. Nucleic Acids Research 2016;44:D184–9.

Craigen WJ, Caskey CT. Expression of peptide chain release factor 2 requires high-efficiency frameshift. Nature 1986;322:273–5.

Craigen WJ, Cook RG, Tate WP et al. Bacterial peptide chain release factors: conserved primary structure and possible frameshift regulation of release factor 2. Proc Natl Acad Sci USA 1985;82:3616–20.

Curran JF, Yarus M. Use of tRNA suppressors to probe regulation of *Escherichia coli* release factor 2. J Mol Biol 1988;203:75–83.

Das G, Thotala DK, Kapoor S et al. Role of 16S ribosomal RNA methylations in translation initiation in Escherichia coli. EMBO J 2008;27:840–51.

Deatherage DE, Barrick JE. Identification of mutations in laboratory-evolved microbes from next-generation sequencing data using breseq. Methods Mol Biol 2014;1151:165–88.

Devaraj A, Shoji S, Holbrook ED et al. A role for the 30S subunit E site in maintenance of the translational reading frame. RNA 2009;15:255–65.

Diop D, Chauvin C, Jean-Jean O. Aminoglycosides and other factors promoting stop codon readthrough in human cells. Comptes Rendus Biologies 2006;330:71–9.

Donly BC, Edgar CD, Adamski FM et al. Frameshift autoregulation in the gene for *Escherichia coli* release factor 2: partly functional mutants result in frameshift enhancement. Nucl Acids Res 1990;18:6517–22.

Fan Y, Evans CR, Barber KW et al. Heterogeneity of Stop Codon Readthrough in Single Bacterial Cells and Implications for Population Fitness. Molecular Cell 2017;67:826–836.e5.

Farr AD, Vasileiou C, Lind PA et al. An extreme mutational hotspot in nlpD depends on transcriptional induction of rpoS. PLoS Genet 2025;21:e1011572.

Freistroffer DV, Kwiatkowski M, Buckingham RH et al. The accuracy of codon recognition by polypeptide release factors. Proc Natl Acad Sci USA 2000;97:2046–51.

Helser TL, Davies JE, Dahlberg JE. Mechanism of Kasugamycin Resistance in Escherichia coli. Nature New Biology 1972;235:6–9.

Herbst KL, Nichols LM, Gesteland RF et al. A mutation in ribosomal protein L9 affects ribosomal hopping during translation of gene 60 from bacteriophage T4. Proc Natl Acad Sci USA 1994;91:12525–9.

Herr AJ, Nelson CC, Wills NM et al. Analysis of the roles of tRNA structure, ribosomal protein L9, and the bacteriophage T4 gene 60 bypassing signals during ribosome slippage on mRNA. Journal of Molecular Biology 2001;309:1029–48.

Higashi K, Kashiwagi K, Taniguchi S et al. Enhancement of +1 frameshift by polyamines during translation of polypeptide release factor 2 in *Escherichia coli*. J Biol Chem 2006;281:9527–37.

Ho AT, Hurst LD. Variation in Release Factor Abundance Is Not Needed to Explain Trends in Bacterial Stop Codon Usage. Molecular Biology and Evolution 2022;39:msab326.

Johnson DBF, Xu J, Shen Z et al. RF1 knockout allows ribosomal incorporation of unnatural amino acids at multiple sites. Nat Chem Biol 2011;7:779–86.

Jørgensen F, Adamski FM, Tate WP et al. Release Factor-dependent False Stops are Infrequent in Escherichia coli. Journal of Molecular Biology 1993;230:41–50.

Khomarbaghi Z, Ngan WY, Ayan GB et al. Large-scale duplication events underpin population-level flexibility in tRNA gene copy number in *Pseudomonas fluorescens* SBW25. Nucleic Acids Research 2024;52:2446–62.

Kim J, Na S-I, Kim D et al. UBCG2: Up-to-date bacterial core genes and pipeline for phylogenomic analysis. J Microbiol 2021;59:609–15.

Kimura S, Suzuki T. Fine-tuning of the ribosomal decoding center by conserved methyl-modifications in the Escherichia coli 16S rRNA. Nucleic Acids Research 2010;38:1341–52.

King EO, Ward MK, Raney DE. Two simple media for the demonstration of pyocyanin and fluorescin. J Lab Clin Med 1954.

Lalanne J, Parker DJ, Li G. Spurious regulatory connections dictate the expression-fitness landscape of translation factors. Mol Syst Biol 2021;17.

Levinson G, Gutman GA. Slipped-strand mispairing: a major mechanism for DNA sequence evolution. Molecular Biology and Evolution 1987.

Loughran G, Li X, O’Loughlin S et al. Monitoring translation in all reading frames downstream of weak stop codons provides mechanistic insights into the impact of nucleotide and cellular contexts. Nucl Acids Res 2023;51:304–14.

Malik V, Rodino-Klapac LR, Viollet L et al. Aminoglycoside-induced mutation suppression (stop codon readthrough) as a therapeutic strategy for Duchenne muscular dystrophy. Ther Adv Neurol Disord 2010;3:379–89.

Márquez V, Wilson DN, Tate WP et al. Maintaining the ribosomal reading frame. Cell 2004;118:45–55.

McNutt ZA, Gandhi MD, Shatoff EA et al. Comparative analysis of anti-Shine-Dalgarno function in *Flavobacterium johnsoniae* and *Escherichia coli*. Front Mol Biosci 2021;8:787388.

Mikuni O, Kawakami K, Nakamura Y. Sequence and functional analysis of mutations in the gene encoding peptide-chain-release factor 2 of *Escherichia coli*. Biochimie 1991;73:1509–16.

Müller MBD, Becker T, Denk T et al. The ribosome as a platform to coordinate mRNA decay. Nucleic Acids Research 2025;53:gkaf049.

Naeem FM, Gemler BT, McNutt ZA et al. Analysis of programmed frameshifting during translation of prfB in Flavobacterium johnsoniae. RNA 2023:rna.079721.123.

Olm MR, Brown CT, Brooks B et al. dRep: a tool for fast and accurate genomic comparisons that enables improved genome recovery from metagenomes through de-replication. The ISME Journal 2017;11:2864–8.

Pavlov MYu, Freistroffer DV, Dincbas V et al. A direct estimation of the context effect on the efficiency of termination. J Mol Biol 1998;284:579–90.

Persson BC, Atkins JF. Does disparate occurrence of autoregulatory programmed frameshifting in decoding the release factor 2 gene reflect an ancient origin with loss in independent lineages? J Bacteriol 1998;180:3462–6.

Pletnev P, Guseva E, Zanina A et al. Comprehensive Functional Analysis of Escherichia coli Ribosomal RNA Methyltransferases. Front Genet 2020;11:97.

Poole ES, Brown CM, Tate WP. The identity of the base following the stop codon determines the efficiency of *in vivo* translational termination in *Escherichia coli*. EMBO J 1995;14:151–8.

Prince CR, Lin IN, Feaga HA. Conservation and evolution of the programmed ribosomal frameshift in *prfB* across the bacterial domain. mBio 2025;16:e01055–25.

Raval PK, Ngan WY, Gallie J et al. The layered costs and benefits of translational redundancy. eLife 2023;12:e81005.

Régnier P, Arraiano CM. Degradation of mRNA in bacteria: emergence of ubiquitous features. Bioessays 2000;22:235–44.

Rengby O, Arnér ESJ. Titration and conditional knockdown of the *prfB* gene in *Escherichia coli*: Effects on growth and overproduction of the recombinant mammalian selenoprotein thioredoxin reductase. Appl Environ Microbiol 2007;73:432–41.

Riegger RJ, Caliskan N. Thinking outside the frame: Impacting genomes capacity by programmed ribosomal frameshifting. Front Mol Biosci 2022;9:842261.

Sanders CL, Curran JF. Genetic analysis of the E site during RF2 programmed frameshifting. RNA 2007;13:1483–91.

Sharma H, Anand B. Ribosome assembly defects subvert initiation Factor3 mediated scrutiny of bona fide start signal. Nucleic Acids Research 2019;47:11368–86.

Silby MW, Cerdeño-Tárraga AM, Vernikos GS et al. Genomic and genetic analyses of diversity and plant interactions of Pseudomonas fluorescens. Genome Biol 2009;10:R51.

de Silva E, Krishnan J, Betney R et al. A mathematical modelling framework for elucidating the role of feedback control in translation termination. J Theor Biol 2010;264:808–21.

Smith AM, Costello MS, Kettring AH et al. Ribosome collisions alter frameshifting at translational reprogramming motifs in bacterial mRNAs. Proc Natl Acad Sci USA 2019;116:21769–79.

Van Buul CPJJ, Visser W, Van Knippenberg PH. Increased translational fidelity caused by the antibiotic kasugamycin and ribosomal ambiguity in mutants harbouring the *ksgA* gene. FEBS Letters 1984;177:119–24.

Wei Y, Wang J, Xia X. Coevolution between Stop Codon Usage and Release Factors in Bacterial Species. Mol Biol Evol 2016;33:2357–67.

Wei Y, Xia X. The role of +4U as an extended translation termination signal in bacteria. Genetics 2017;205:539–49.

Weiss RB, Dunn DM, Atkins JF et al. Slippery runs, shifty stops, backward steps, and forward hops: - 2, -1, +1, +2, +5, and +6 ribosomal frameshifting. Cold Spring Harb Symp Quant Biol 1987;52:687–93.

Weiss RB, Dunn DM, Dahlberg AE et al. Reading frame switch caused by base-pair formation between the 3′ end of 16S rRNA and the mRNA during elongation of protein synthesis in *Escherichia coli*. EMBO J 1988;7:1503–7.

Williams JM, Donly BC, Brown CM et al. Frameshifting in the synthesis of *Escherichia coli* polypeptide chain release factor two on eukaryotic ribosomes. European Journal of Biochemistry 1989;186:515–21.

Wong T-Y, Fernandes S, Sankhon N et al. Role of premature stop codons in bacterial evolution. J Bacteriol 2008;190:6718–25.

Worth CL, Gong S, Blundell TL. Structural and functional constraints in the evolution of protein families. Nat Rev Mol Cell Biol 2009;10:709–20.

Zhang H, Lyu Z, Fan Y et al. Metabolic stress promotes stop codon readthrough and phenotypic heterogeneity. Proc Natl Acad Sci USA 2020;117:22167–72.

Zhang X-X, Rainey PB. Construction and validation of a neutrally-marked strain of Pseudomonas fluorescens SBW25. Journal of Microbiological Methods 2007;71:78–81.

Zhang Y, Nersisyan L, Fürst E et al. Ribosomes modulate transcriptome abundance via generalized frameshift and out-of-frame mRNA decay. Molecular Cell 2025;85:2017–2031.e7.

