## Supplementary Figures S1 to S4 and Supplementary Methods for "Compensatory evolution facilitates loss of *prfB* autoregulation in *Pseudomonas fluorescens* SBW25"

Sungbin Lim, Frederic Bertels, Javier Lopez-Garrido* and Jenna Gallie*†
Max Planck Institute for Evolutionary Biology, 24306 Plön, Germany

†Current address: Cellora, Pōkeno 2402, New Zealand

Content:

- Supplementary Figures S1 to S4
- Supplementary Methods

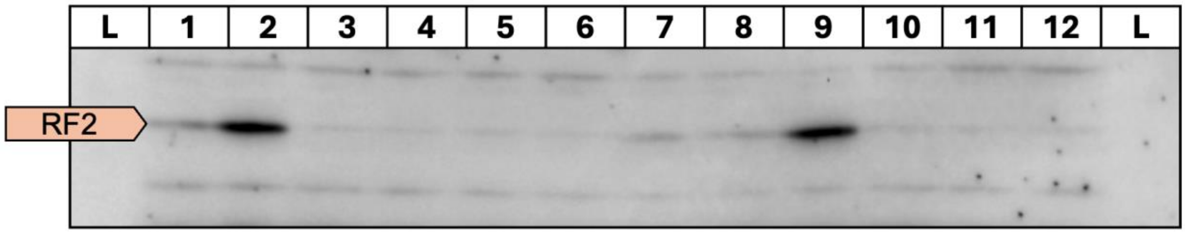


**Supplementary Figure S1. Western blot analysis of RF2 in wild-type *Pseudomonas fluorescens* SBW25 and in different mutant strains.** The RF2 band (expected size: 41 kDa) is indicated by an arrow box. Nonspecific bands are also observed above and below the RF2 band. The image was cropped to display target region. The image was taken with a 300-second exposure. Full images including alternative exposure times are provided in the Supplementary Data. Lane annotations are as follows:

L – Protein ladder

1 – *prfB*$^{\text{WT}}$

2 – *prfB*$^{\Delta\text{Stop}}$

3 – *prfB*$^{\text{CUA}}$

4 – *prfB*$^{\text{mSD}}$

5 – *prfB*$^{\text{CUA}}$ Δ*rsmA*

6 – *prfB*$^{\text{CUA}}$ Δ*rplI*

7 – Δ*rsmA*

8 – Δ*rplI*

9 – *prfB*$^{\text{CUA}}$ Iso-PRF-1

10 –*prfB*$^{\text{CUA}}$ Iso-*rsmA*-4a

11 –*prfB*$^{\text{CUA}}$ Iso-*rsmH*

12 –*prfB*$^{\text{CUA}}$ Iso-*rplI*

L – Protein ladder (duplicate)

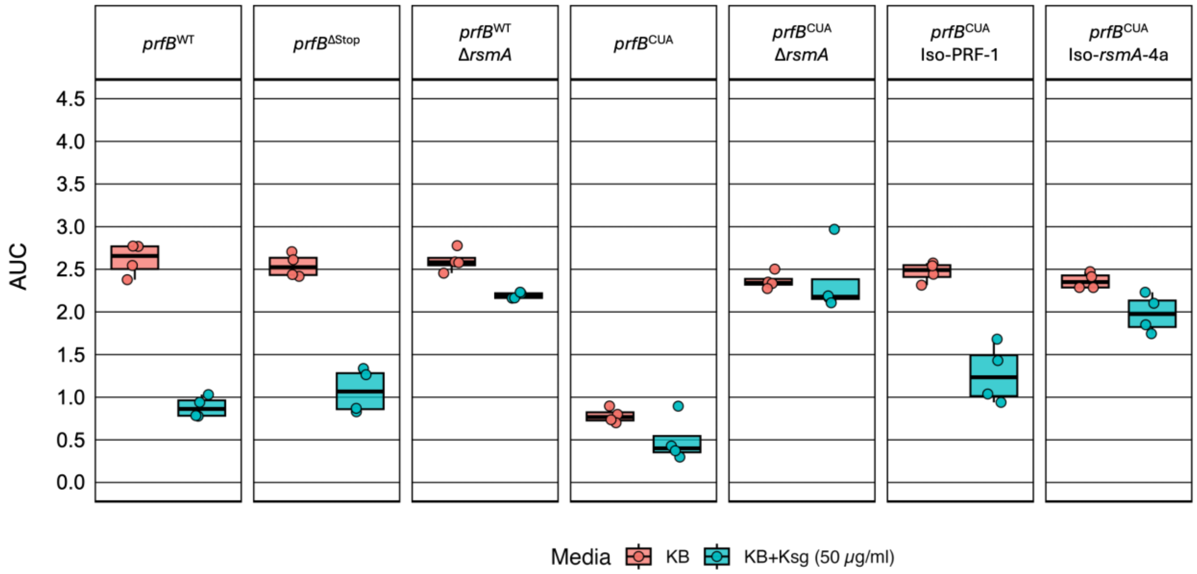


**Supplementary Figure S2. Impact of mutations in *prfB* and *rsmA* on kasugamycin resistance.** Area under the growth curve (AUC) during 12 hours of growth with the indicated *P. fluorescens* strains (top of each panel) in KB medium in the absence (red) or in the presence (green) of 50 µg/ml kasugamycin (Ksg). Representative evolved isolates were included in the analysis. Data represents four independent replicates. See the legend of Figure 2C for box plot description.

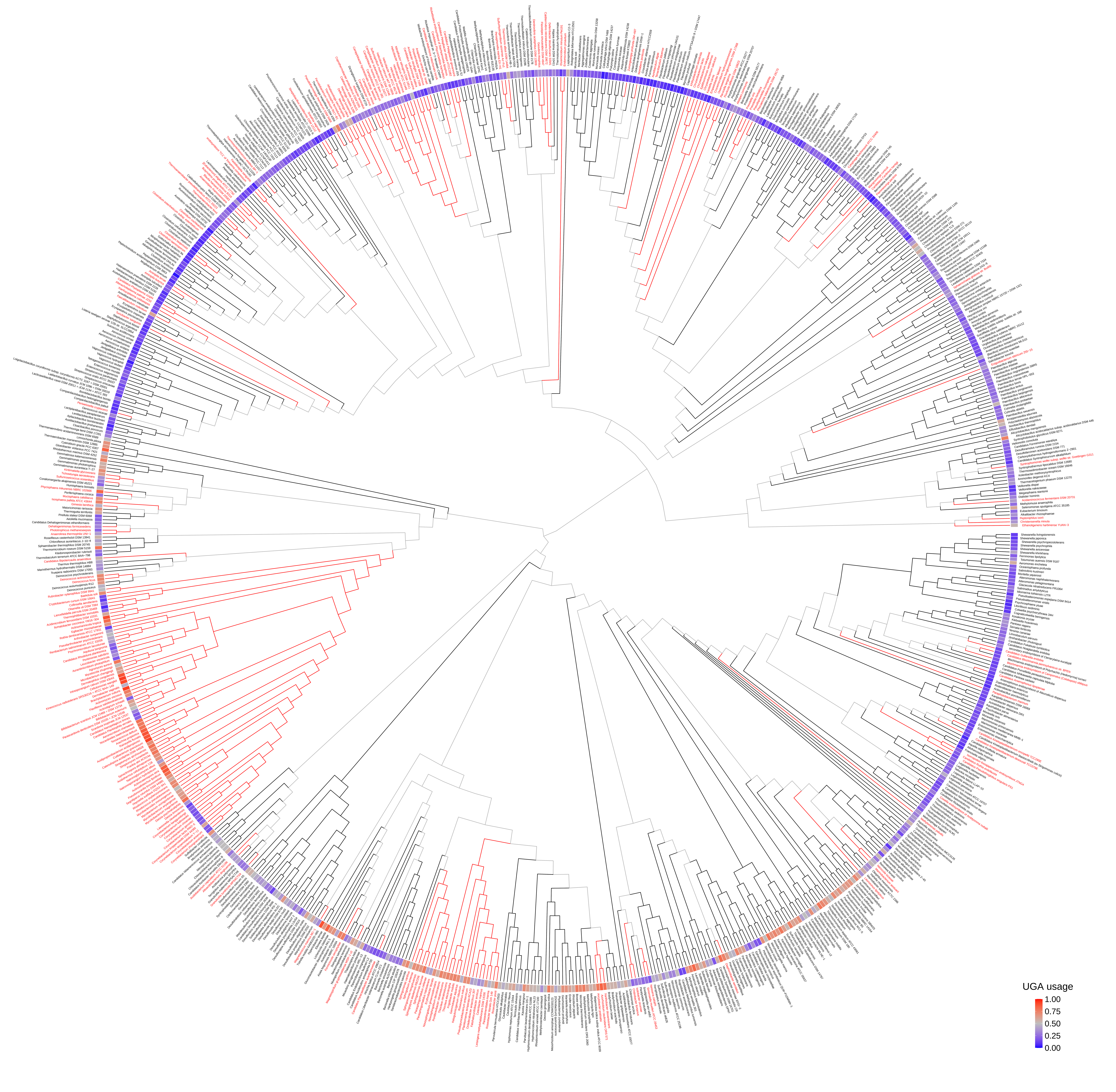
UGA usage
1.00
0.75
0.50
0.25
0.00

**Supplementary Figure S3. High-resolution phylogenetic tree based on core gene sequence with species name indicated** (A) Phylogenetic tree of *prfB* (818 species) with high-resolution including species name on the tip of each node. Leaves and branches for $PRF^+$ species are black; $PRF^-$ species are red. Species-level UGA stop codon usage is represented as a color-coded heatmap, with a frequency of 0.5 shown in gray, higher frequencies in red, and lower frequencies in blue. The tree was built with UBCG2 (Kim *et al.* 2021).

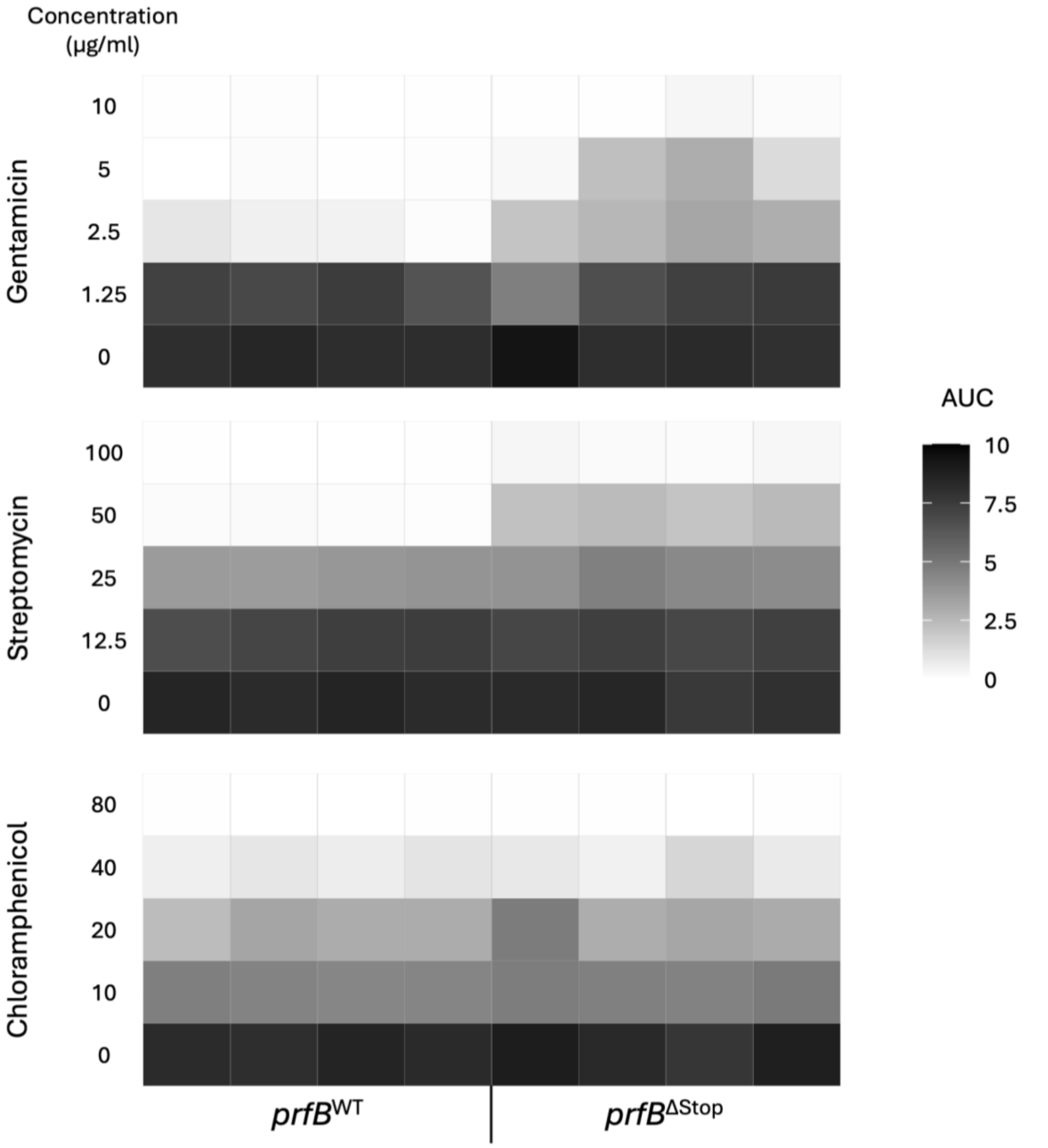


**Supplementary Figure S4. Impact of deletion of autoregulation in *prfB* on antibiotic resistance.** Area under the growth curve (AUC) during 24 hours of growth with the indicated *P. fluorescens* strains (bottom of each panel) in KB medium with different antibiotics at the indicated concentrations. Heatmaps of four replicates are shown per strain and antibiotic concentration.

**Supplementary Methods. Details about different experimental evolution runs**

Three independent evolution experiments were performed (Exp1 to Exp3). The specific experimental setup for each experiment was as follows:

Exp1

- Lineage composition: 8 independent lineages of SBW25-*lacZ* and 8 independent lineages of the *prfB*$^{CUA}$ mutant.
- Transfer volume: 4 lineages of each genotype were transferred with 4 µl (1:1000 dilution), and the other 4 lineages were transferred with 40 µl (1:100 dilution).
- Excluded lineages: One *prfB*$^{CUA}$ lineage was excluded due to contamination.
- All isolates were selected at Week 1.

Exp2

- Lineage composition: 1 lineage of SBW25-*lacZ* and 16 independent lineages of the *prfB*$^{CUA}$ mutant.
- Transfer volume: 40 µl for all lineages (1:100 dilution).
- Excluded lineages: One *prfB*$^{CUA}$ lineage was excluded due to contamination.
- All isolates were selected at Week 1.

Exp3

- Lineage composition: 2 independent lineages of SBW25-*lacZ*, 16 independent lineages of the *prfB*$^{CUA}$ mutant.
- Transfer volume: 40 µl for all lineages (1:100 dilution).
- Excluded lineages: Six *prfB*$^{CUA}$ lineages were excluded due to contamination.
- In one lineage, a single isolate, Iso-rsmA-10, was isolated on Week 1, and another isolate, Iso-PRF-2, was isolated on Week 3. All other isolates were acquired separately from each lineages in Week 1.

The table below indicates the number of isolates obtained from the different experimental runs:

| Gene/Allele ID | Exp1 | Exp2 | Exp3 | Total |
|---|---|---|---|---|
| *prfB* / Iso-PRF-1 | 2 | | | 2 |
| *prfB* / Iso-PRF-2 | 1 | | 1 | 2 |
| *rsmA* / Iso-*rsmA*-1a | 1 | 1 | | 2 |
| *rsmA* / Iso-*rsmA*-1b | | | 1 | 1 |
| *rsmA* / Iso-*rsmA*-2a | | 1 | | 1 |
| *rsmA* / Iso-*rsmA*-2b | 1 | | | 1 |
| *rsmA* / Iso-*rsmA*-3 | | 1 | | 1 |
| *rsmA* / Iso-*rsmA*-4a | 1 | 9 | 4 | 14 |
| *rsmA* / Iso-*rsmA*-4b | | | 1 | 1 |
| *rsmA* / Iso-*rsmA*-5 | 1 | | | 1 |
| *rsmA* / Iso-*rsmA*-6 | | 1 | | 1 |
| *rsmA* / Iso-*rsmA*-7 | | 1 | | 1 |
| *rsmA* / Iso-*rsmA*-8 | | | 1 | 1 |
| *rsmA* / Iso-*rsmA*-9 | | | 1 | 1 |
| *rsmA* / Iso-*rsmA*-10 | | | 2 | 2 |
| *rsmH* / Iso-*rsmH* | | 1 | | 1 |
| *rplI* / Iso-*rplI* | | 1 | | 1 |
| Total | 7 | 16 | 11 | 34 |

Kim J, Na S-I, Kim D *et al.* UBCG2: Up-to-date bacterial core genes and pipeline for phylogenomic analysis. *J Microbiol* 2021;**59**:609–15.
